# Design trade-offs and robust architectures for combined transcriptional and translational resource allocation controllers

**DOI:** 10.1101/2020.02.11.944215

**Authors:** Alexander P.S. Darlington, Declan G. Bates

## Abstract

Recent work on engineering synthetic cellular circuitry has shown that non-regulatory interactions brought about through competition for shared gene expression resources, such as RNA polymerase and ribosomes, can result in degraded performance or even circuit failure. Transcriptional and translational resource allocation controllers based on orthogonal ‘circuit-specific’ gene expression machineries have previously been separately designed to enforce modularity and improve circuit performance. Here we investigate the potential advantages, challenges, and design trade-offs involved in combining transcriptional and translational resource allocation into one overarching centralised control system. We design a number of biologically feasible controllers that reduce coupling at both the transcriptional and translational levels simultaneously, and identify some key performance tradeoffs. We apply tools from robust control theory to rigorously quantify the impact of uncertainty/variability arising due to experimental implementations on the operation of such controllers. Based on these results, we identify promising architectures for the construction of robust dual transcriptional–translational resource allocation controllers.

## 1 Introduction

Synthetic gene circuits can be designed to perform complex computations and information processing in living cells with applications in biomedicine, chemistry and environmental sciences [1]. By introducing synthetic gene circuits into microbial hosts, synthetic biologists and biotechnologists are able to control cell function. However, often these initial designs fail due to the effect of unforeseen interactions between the circuit and host cell or due to host constraints [2, 3]. In addition, circuits produced in one strain can behave both quantitatively and qualitatively differently when transferred to other strains, due to the subtle impact of changing genetic context [4, 5, 6, 7].

A key cause of context dependent dysfunction of gene circuits is the competition for shared gene expression resources, such as host RNA polymerases and ribosomes [8, 7, 9, 10]. Two independently characterised modules, when brought together in a synthetic circuit, interact though the use of common resource pools; e.g. as one gene is induced it indirectly inhibits other genes by sequestering finite cellular resources. This impact is greater for protein encoding genes; where the induction of the second gene results in both a transcriptional and translational disturbance as RNA polymerase molecules are sequestered for mRNA production and ribosomes are sequestered for protein production. Transcriptional disturbances, caused by the induction of non-protein encoding small RNAs, can also result in perturbations at translational levels, due to competition for RNA polymerase. In *E. coli*, transcriptional coupling often results in more subtle effects than translation coupling [8, 11], but has been shown to have more significant effects in mammalian systems [12, 13].

Circuit-specific ‘orthogonal’ gene expression resources which only transcribe/translate circuit genes (and not host genes) have been proposed to alleviate some aspects of host circuit interactions in *E. coli* [14]. These systems utilise a bacteriophage RNA polymerase for transcription and synthetic ribosomal RNAs to render host ribosomes orthogonal. It has been proposed that these orthogonal gene expression resources can form a cellular ‘virtual machine’ [15] and allow the decoupling of circuit genes without extensive re-design [18].

At the transcriptional level, Kushwaha and Salis have previously designed and implemented a ‘universal bacterial expression resource’ (UBER) to alleviate the impact of species-context dependency, [6]. This system is based on an RNA polymerase from the T7 bacteriophage, which forms an orthogonal transcriptional resource and is able to transcribe mRNAs in a range of different species (Figure 1A). The orthogonal RNAP (o-RNAP) transcribes its own mRNA creating a positive feedback loop. However, at high concentrations, the T7 RNAP is toxic and results in reduced cell growth. Therefore, the o-RNAP transcription must be tightly controlled using negative feedback, by placing it under the regulation of the repressor TetR, which itself is transcribed by the o-RNAP. In [16], Segall-Shapiro *et al.* developed a set of compatible artificial fragments of T7 RNA polymerase. When co-expressed these fragments bind to form a functional RNA polymerase. They used these components to develop a ‘transcriptional resource allocator’ which buffers the impact of different gene copy numbers. The core component of RNA polymerase *β* is constitutively expressed but only becomes functional when bound by the α-fragment which is co-expressed with the circuit genes (Figure 1D).

**Figure 1:**
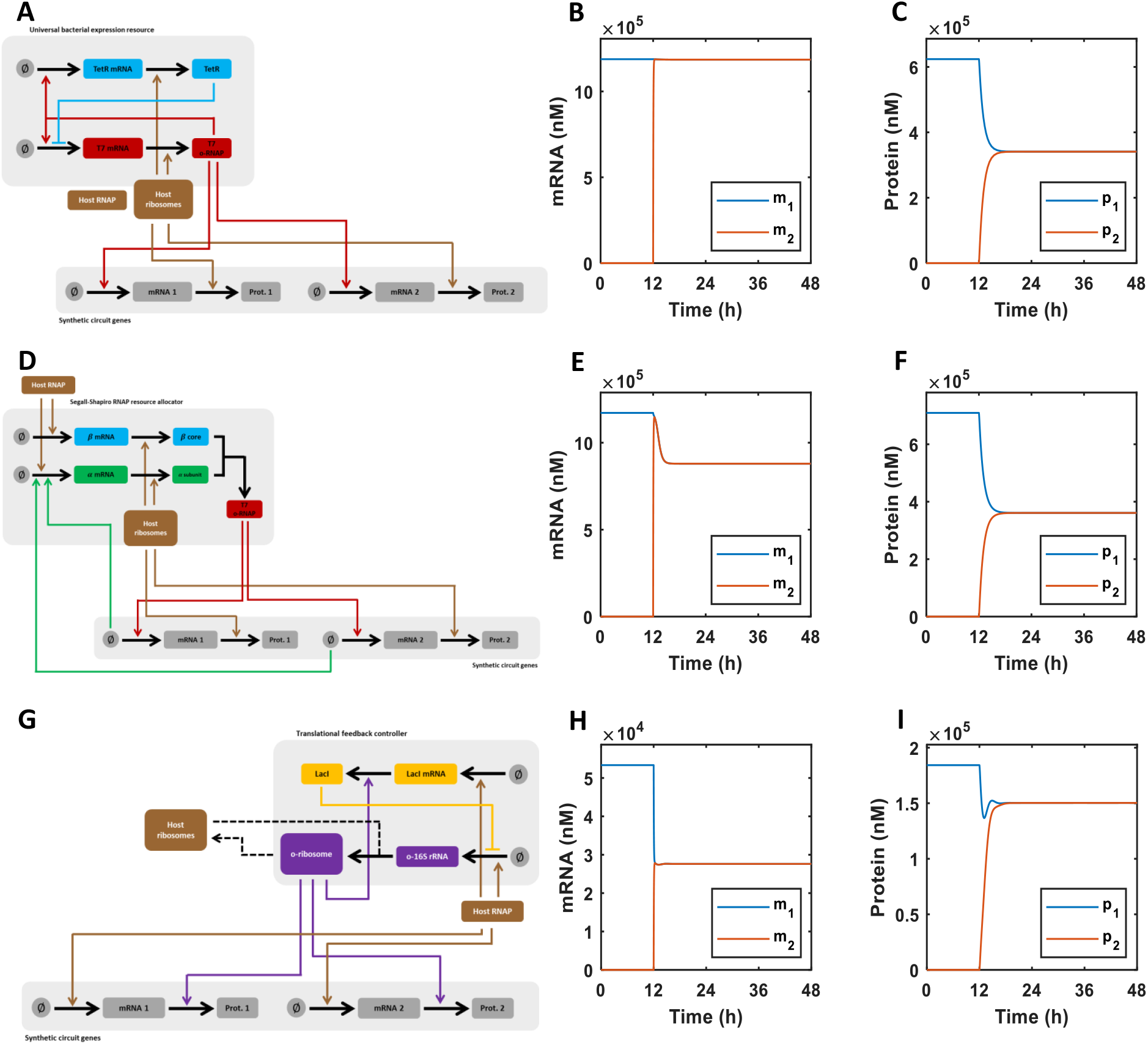
Gene decoupling by transcriptional and translational controllers. The behaviour of candidate transcriptional and translational control systems in isolation. We simulated the ability of each prototype control system to decouple co-expressed genes. We consider the impact on a constitutively expressed gene (*g*_1_) of the induction of a second gene (*g*_2_) at 12 h. This results in a transcriptional disturbance (as promoters compete for RNA polymerase) *and* a translational disturbance (as mRNAs compete for ribosomes). **(A)** Architecture of the universal bacterial expression resource (UBER) which supplies orthogonal RNA polymerases to the circuit. **(B)** The UBER controller successfully mitigates the transcriptional disturbance applied at 12 h. **(C)** The UBER controller is unable to mitigate the disturbance at the translational level. **(D)** Architecture of the fragmented RNA polymerase resource allocation controller (FRAG) which supplies orthogonal RNA polymerases to the circuit. **(E)** The fragmented RNA polymerase controller is able to mitigate the transcriptional disturbance at 12 h to some extent with the fall in *m*_1_ only being 25% rather than 50%. **(F)** The FRAG controller is unable to mitigate the disturbance at the translational level. **(G)** Architecture of the orthogonal ribosome-based translational controller (OR) dynamically supplies translational activity to the circuit. **(H)** The translational controller has no impact on the transcriptional disturbance with *m*_1_ falling by 50% upon activation of the second protein-encoding gene. **(I)** The translational controller decouples genes at the translational level reducing the fall in *p*_1_ from 50% to only 20%.

To alleviate translational resource competition, we have previously designed and implemented a translational resource allocation controller in *E. coli* [17, 18]. The ribosome is a large ribonucleoprotein complex encoded by multiple rRNA and r-protein genes; therefore, unlike in the case of transcription, there does not exist a truly orthogonal ribosome which can be co-opted from another species. However, quasi-orthogonal ribosomes (o-ribosomes) can be created using synthetic 16S rRNAs which target the ribosome machinery to mRNAs that contain the complementary ribosome binding site (RBS) sequence (e.g. [14]). The translational controller works by regulating the production of the synthetic 16S rRNA. This negative feedback controller takes the form of a repressor protein (LacI was used in [17]) which inhibits 16S rRNA production and itself is translated by the o-ribosome pool. As demand for orthogonal translation increases, the level of the repressor falls due to resource competition and so o-rRNA, and hence orthogonal ribosome, production increases (matching demand).

In this paper, we consider how orthogonal transcriptional and translational resource allocation systems can be combined to function as dual transcription–translation controllers that decouple genes at both levels of expression. We show that interactions between separately functional transcriptional and translational controllers can result in instability when they are implemented simultaneously. We design dual controllers that overcome this problem, and identify some fundamental trade-offs between decoupling performance and gene expression levels. Finally, we demonstrate how analytical tools from Robust Control Theory can be used to rigorously quantify the robustness of different controller architectures and therefore guide selection of designs for future biological implementation.

## 2 Results

### 2.1 Modelling resource-mediated coupling at the transcriptional and translational levels demonstrates the need for combined resource allocation

Initially we developed a simple ordinary differential equation model taking into account transcription (modelled as RNA polymerase binding/unbinding to/from a promoter and mRNA birth), translation (ribosome binding/unbinding to/from an mRNA and protein birth) and dilution of all species and intermediate complexes. This base model takes account of usage of host gene expression resources and therefore captures how resource limitations create non-regulatory couplings. For full model details see Methods (Section 4.1). We next augment the base model with the additional orthogonal gene expression resource and its respective control system, changing the transcriptional or translational apparatus from host to orthogonal, as appropriate.

The universal bacterial expression resources developed in [6] decouples co-expressed genes at the transcriptional level with the mRNA of gene 1, *m*_1_, showing no disturbance upon the activation of gene 2 (Figure 1B). However, this does not propagate to the translational level due to competition for translational resources (Figure 1C). The transcriptional controller developed in [16], shows poorer performance with significant coupling remaining when the transcriptional disturbance is applied: the mRNA of gene 1 falls 25% when gene 2 is induced at 12 h (Figure 1G). This still represents an improvement from a 50% fall in gene 1 in absence of control (Figure S1A). Again this decoupling does not propagate to the translational level with protein 1 falling 50% upon induction of protein 2. (Figure 1F). Both controller topologies achieve their transcriptional decoupling action by increasing the effective concentration of free RNA polymerase, rather than increasing orthogonal RNA polymerase production to match demand (Figure S1). In the absence of control, increasing the concentration of RNAP (either host or orthogonal) *in silico* results in decreased mRNA coupling (Figure S1E). The poorer performance of the Segall-Shapiro *et al.* controller is due to a fall in RNA polymerase concentration due to *translational* coupling between the polymerase expression and circuit genes - this moves the system from a regime of low coupling to moderate coupling (Figure S1E). A significant number of the universal bacterial expression resource controllers (approximately 40%) show decoupling at the transcriptional and translational levels. However, these controllers have negligible output, with the designs corresponding to those with high expression of *p_q_*. This results in negligible production of the orthogonal RNA polymerase, resulting in low mRNA levels and reduced translational competition. A small number (8%) of designs with this controller architecture also showed unstable oscillatory behaviour.

In [18], we fully analysed a resource allocation controller at the translational level (in the absence of RNAP competition) to derive design guidelines for potential biological implementations. Applying these findings, we simulated a controller composed of the tightly binding multimeric repressor (here LacI), high o-rRNA copy number and low transcription factor copy number (Figure 1G). In the presence of RNA polymerase competition, this controller is not perfect but still reduces the fall in protein 1 when protein 2 is induced to 20% (Figure 1I). Decoupling is not complete (i.e. no fall in *p*_1_ upon addition of *p*_2_) due to the presence of coupling at the transcriptional level (Figure 1H). Simulating the response of the controller to an RNA-only input (that is a transcriptional but *not* a translational disturbance) demonstrates that the controller is not able to respond to transcriptional competition; i.e. there is a 10% decrease in *p***1** even though there is no ribosomal competition, due to competition for the host RNAP (Fig. S2F).

This preliminary analysis clearly highlights the need to implement resource allocation at both transcriptional and translational levels simultaneously in order to deal with all possible non-regulatory coupling arising from host-circuit competition for finite cellular resorces.

### 2.2 Dual transcriptional-translational resource allocation controllers: potential architectures and design tradeoffs

We now consider the ability of the transcriptional (UBER or FRAG) and translational controllers (OR) to operate together simultaneously to decouple circuits at both the transcriptional and translational levels, i.e. to reject both transcriptional disturbances caused by mRNA or sRNA expression and translational disturbances caused by mRNA expression. This was achieved by setting both gene expression resources for the circuit genes to their orthogonal counterparts (Figure 2). We name these novel dual control systems UBER-OR and FRAG-OR based on how the orthogonal RNA polymerase activity is controlled. Both control systems utilise the same translational resource allocation controller (OR). We do not change each controller’s own gene expression resource usage and so each controller’s internal topology remains the same. Combining the putative transcriptional and translational controllers (which function well in isolation - see Figure 1), into two potential ‘dual’ controllers results in the emergence of instability (Figure S3A, B) or poor performance at the translational level (Figure S3B, D).

**Figure 2:**
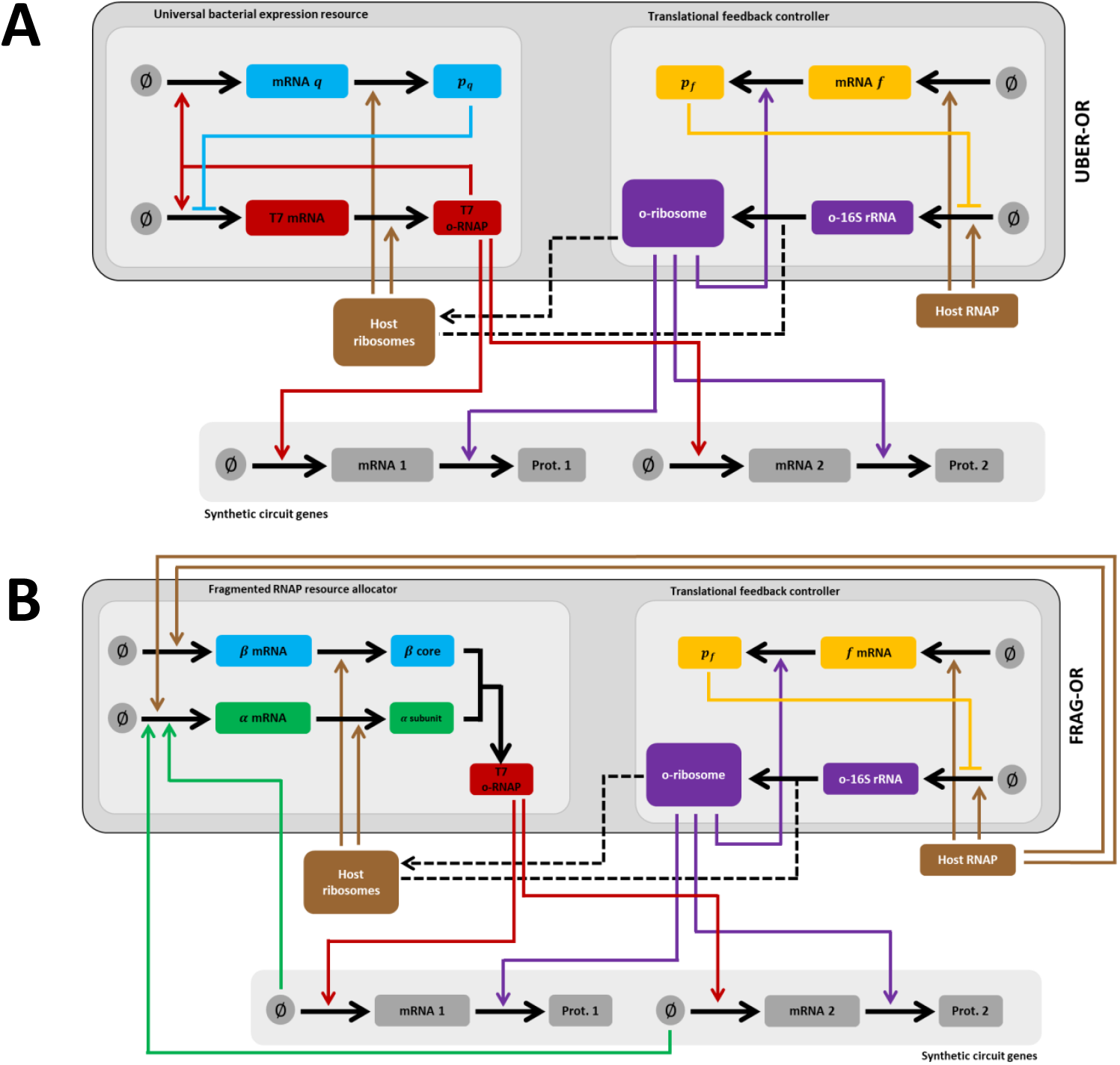
Architecture of the combined dual transcriptional-translational controllers. **(A)** UBER-OR is based on the universal bacterial expression resource developed in [6] and the translational controller developed in [17]. **(B)** FRAG-OR is based on the fragmented RNA polymerase resource allocator developed in [16] and the translational controller developed in [17].

To understand better the operation of these two dual controllers we analysed a number of experimentally feasible designs. We created a discrete, rather than continuous, design space across a range of experimentally tunable parameters representing promoter and ribosome binding site strengths, controller gene copy numbers, and transcription factors. The choice of transcription factor impacts *α_r,x_* representing the dissociation constant, *η_x_* the transcription multimeric state and the dissociation constant of the target promoter (*ξ_f,x_*) (where *x* represents the species being regulated). These parameters are not independently designable and so we simulate the action of the common repressors tetR, lacI and cI. (Note that the UBER-OR requires the use of two transcription factors as resource controller proteins which we do not allow to be the same, i.e. *p_q_* = *p_f_*).

For each controller architecture and numerical design, we discarded controllers which demonstrate instability (e.g. oscillatory behaviour) and poor output performance. We define performance in three ways: (1) mRNA coupling (i.e. the change in *m*_1_ when *g*_2_ is activated), (2) protein coupling (i.e. the change in *p*_1_ when *g*_2_ is induced) and (3) final protein concentration. We collapse metrics (1) and (2) into one. The ideal dual controller would perfectly decouple genes at the mRNA and protein levels - i.e. *m*_1_ and *p*_1_ would not change (equivalent to (0, 0) in a two dimensional performance space). First we scaled each coupling metric by the maximum absolute coupling and then considered the Euclidean distance for each pair of (mRNA coupling, protein coupling) values. We refer to this as the 2D score. We identify the best ‘dual’ controllers using this metric (see Methods). Proteins form the main actuators of synthetic gene circuits and so we also consider final proteins levels as part of our performance assessment.

This analysis reveals that UBER-OR (Figure 2A) gives superior decoupling with the lowest 2D (Figure 3A) in comparison to FRAG-OR (based on the fragmented RNA polymerase) (Figure 2B). However, the latter architecture tends to produce controllers with higher protein production levels (Figure 3B), indicating a tradeoff between these different performance objectives.

**Figure 3:**
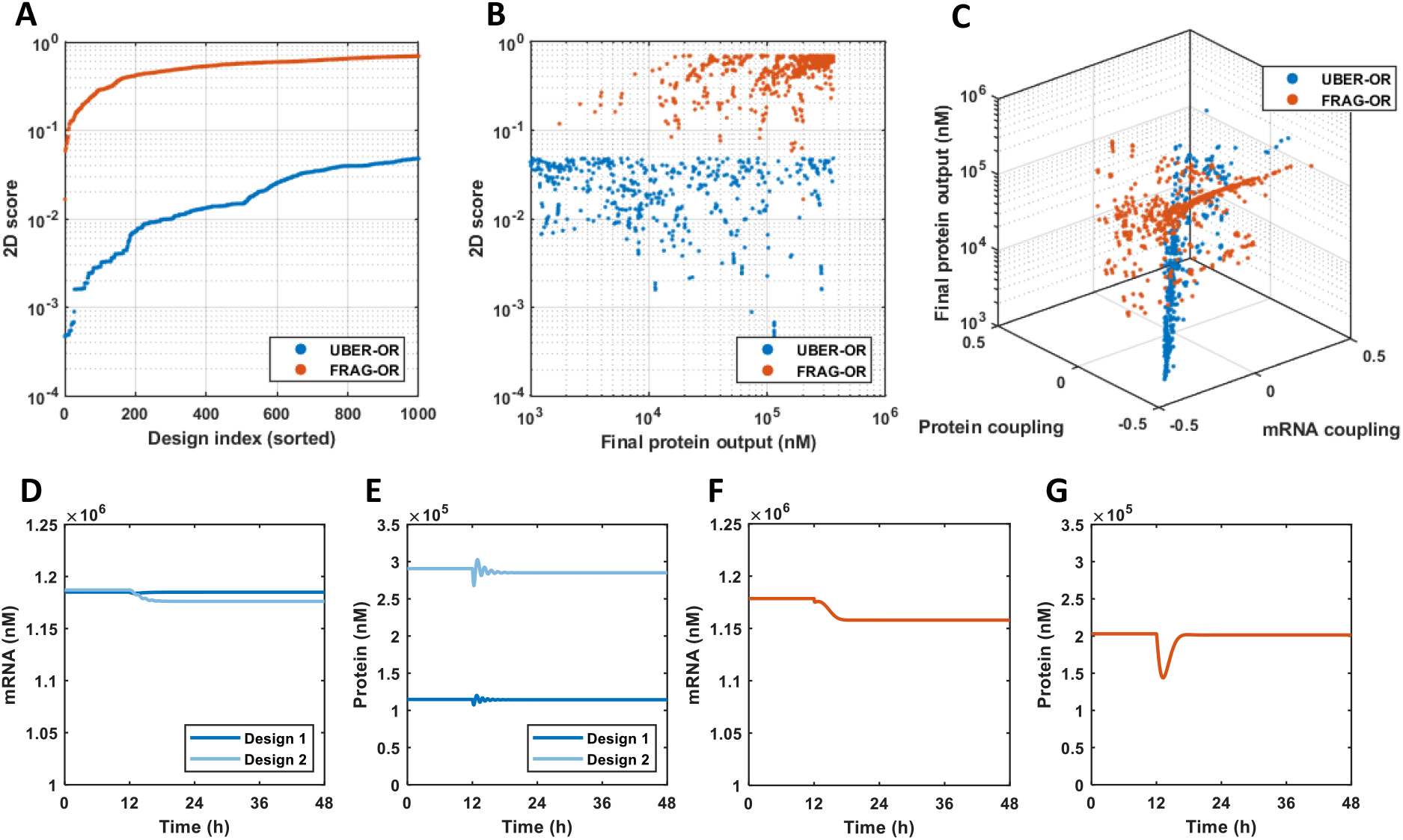
Performance of dual transcriptional–translational controllers. Candidate controllers where simulated as described in the methods. mRNA and protein coupling is calculated as the change in *m*_1_ or *p*_1_ (respectively) upon the induction of the second gene at 12 h. The 2D score is a one dimensional representation of the two dimensional mRNA coupling-protein coupling space and is calculated as outlined in the Methods. **(A)** The two dimensional score of the best 1,000 controllers for each controller architecture. **(B)** Two dimensional score plotted again the final protein concentration for both controller topologies. **(C)** The performance of the best 1,000 controllers in terms of transcriptional and translational coupling and final protein concentration. **(D)** Change in the mRNA concentration in response to the induction of gene 2 of the best design (Design 1) of the combined controller UBER-OR. Design 2 represents a controller with improved gene expression but decreased 2D score. **(E)** Change in the protein concentration in response to the induction of gene 2 of the best design of the combined UBER-OR controller. Design 2 represents a controller with improved gene expression but decreased 2D score. **(F)** Change in the mRNA concentration in response to the induction of gene 2 of the best design of the combined FRAG-OR controller. **(G)** Change in the protein concentration in response to the induction of gene 2 of the best design of the combined FRAG-OR controller.

In general both controllers give access to the same regions of decoupling performance space (Figure 3C). The best designs for each potential controller show similar decoupling abilities at both the transcriptional and translational levels (Figure 3). Note also that some controllers show ‘positive’ coupling i.e. the mRNA or protein of gene 1 *rises* upon induction of gene 2.

Analysing the best decoupling (i.e. lowest 2D score) designs for UBER-OR shows that these controllers can be implemented on various combinations of single and medium (20) multicopy plasmids, with the orthogonal RNA polymerase, o-rRNA and translational controller protein (*p_f_*) being carried on the multicopy plasmid and the transcriptional regulator (*p_q_*) being carried on the lower copy counterpart (Figure S4A). The translational controller protein requires a strong RBS (low *β_f,f_*). The orthogonal RNA polymerase RBS strength (*β_f,p_*) takes numerous intermediate values equivalent to dissociation constants spanning three orders of magnitude (1e4-1e6 nM) with lower values being more common. The transcriptional regulator (*p_q_*) requires a weak ribosome binding site. In nearly all of the best controller designs the transcriptional regulator (*p_q_*) is tetR - a monomeric, relatively weak binding repressor. The translational regulator (*p_f_*) always takes the form of a multimeric strongly binding repressor (with lacI giving better performance over cI).

Analysing the best decoupling designs of FRAG-OR shows that these controllers can be implemented on a single plasmid system with the core RNA polymerase (*β*) fragment and the translational regulator *p_f_* being carried on a medium or high copy plasmid and the orthogonal rRNA gene chromosomally integrated (Figure S4B). Some designs require the o-rRNA to be carried on a medium copy plasmid. Note that the RNA polymerase *α*-fragment is carried on the circuit plasmid in all cases. In general, the *α*-fragment has the weakest RBS of the three protein encoding genes. The *β*-fragment of the RNA polymerase requires a strong to medium RBS while the translational regulator requires a strong RBS. In nearly all of the best performing designs selected the translational regulator (*p_f_*) takes the form of a multimeric strongly binding repressor (with lacI giving better performance over cI).

### 2.3 Robustness to uncertainty in experimental implementations

We have shown that it is possible to design dual transcriptional and translational controllers which satisfy performance criteria in terms of decoupling co-expressed genes and giving satisfactory levels of protein output. We have also shown that these controllers can be composed of biologically reasonable parameters corresponding to obtainable promoter copy numbers and transcription factor dynamics. However, at present the construction of such controllers is complicated by the uncertainty in the kinetic parameters of the available biological ‘parts’, with large potential variations reported for many parts [19]. Often, precise measurements of these parameters are only obtainable from *in vitro* measurements, and how these relate to *in vivo* values is usually unknown. Even when a part with a desired set of kinetics exists with small uncertainty, it is not clear how circuit context effects may impact this level of uncertainty; for example the surrounding DNA sequence may cause subtle changes in binding rates. The causes of uncertainty in biological circuit design have been reviewed extensively in [2] and [20]. To take account of these biological realities, here we assess the robustness of both dual controllers to parametric uncertainty; focusing on the ‘designable’ parameters governing the production rates of the orthogonal RNA polymerase and rRNA and the production rates and action of the other controller proteins (Table 1).

**Table 1:** Full model equations. The equations required to create the full circuit–controller models. The uncertain parameters varied in the robustness analysis are listed.

|  | Open loop | UBER txn | FRAG txn | OR tln |
| --- | --- | --- | --- | --- |
| Circuit | 4–7, 8 |  |  |  |
| o-RNAP | none | 14–22, 23–24 | 29–37, 38–39 | none |
| o-ribosome | none | none | none | 47–54, 55–56 |
| h-RNAP | 9 | 25 | 41 | 57 |
| h-ribosome | 10 | 26 | 42 | 58 |
| Uncertain parameters | none | $\omega_p, \xi_{f,p}, \beta_{f,p}, \beta_{f,q}, \omega_q, \alpha_{r,q}, \eta_q$ | $\omega_c, \beta_{f,c}, \beta_{f,a}$ | $\omega_r, \xi_{f,r}, \omega_f, \beta_{f,f}, \alpha_{r,f}, \eta_f$ |
|  | UBER-OR dual controller |  | FRAG-OR dual controller |  |
| Circuit | 4–7, 8 |  |  |  |
| o-RNAP | 14–22, 23–24 |  | 29–37, 38–40 |  |
| o-ribosome | 47–54, 55–56 |  | 47–54, 55–56 |  |
| h-RNAP | 59 |  | 61 |  |
| h-ribosome | 60 |  | 62 |  |
| Uncertain parameters | $\omega_p, \xi_{f,p}, \beta_{f,p}, \omega_r, \xi_{f,r}, \omega_f, \beta_{f,q}, \omega_q, \beta_{f,f}, \alpha_{r,q}, \eta_q, \alpha_{r,f}, \eta_f$ | | $\omega_c, \beta_{f,c}, \beta_{f,a}, \omega_r, \xi_{f,r}, \omega_f, \beta_{f,f}, \alpha_{r,f}, \eta_f$ | |

In engineering, the field of Robust Control Theory is concerned with addressing the stability and performance of control systems when parameters cannot be estimated, set or designed precisely. The structured singular value (*μ*) provides a method to rigorously compare the robustness of alternative controller designs [21, 22] in the face of uncertainty in multiple parameters. *μ* itself represents the inverse of the maximum level of uncertainty which a feedback control system can tolerate without becoming unstable. Efficient algorithms exist to estimate upper and lower bounds on *μ* for problems which can be expressed as linear fraction transformations (LFTs), i.e. problems for which all known and uncertain parameters can be separated out and connected via a particular feedback structure. For the controllers considered here, however, the degree of complexity/nonlinearity of the closed-loop systems, together with the number of uncertain parameters that impact multiple species and their steady states, makes the generation of LFT representations impossible. Instead, we apply the ‘LFT-free’ *μ* estimation algorithm developed in [23] (and refined in [24]). During this process the system is linearised and then transformed from the time domain into the frequency domain where the system’s outputs are represented as complex numbers. This method reformulates the standard *μ* analysis problem as a geometric one, and uses a probabilistic algorithm to search for the intersection of the real and imaginary parts of the Laplace transformed system in n-dimensional uncertainty space (where *n* is the number of uncertain parameters). This search is repeated over a range of input frequencies, hence generating upper and lower bounds for *μ* in the standard fashion. If, for a given maximum level of uncertainty *δ*_max_, the value of *μ* ≥ 1, then there exists an uncertain parameter combination that destabilises the system, i.e. it is not robust to that level of uncertainty.

Applying this method to to the best performing dual controller design based on the UBER architecture (UBER-OR) reveals a poor level of robustness, with a *μ* estimate of 14.11 < *μ* < 73.8 for an uncertainty level of ±20% in all parameters (Figure 4A). This estimate is based on a linearisation of the model, but the potential instability was confirmed via Monte Carlo simulations of the original non-linear model which identified a number of parameter sets within the allowed level of uncertainty which show oscillatory behaviour (Figure 4B). Repeating this analysis for all of the controller designs for this architecture shows that stability cannot be guaranteed for any biologically reasonable level of uncertainty - the *μ*-upper bound is greater than or equal to 1 for all *δ* > 10% (Figure S5B). In fact, the *μ* analysis guarantees *instability* for uncertainty levels of 10% or greater for 78 of the 100 best controllers (i.e. the lower *μ* bound is also more than 1) (Figure S5A).

**Figure 4:**
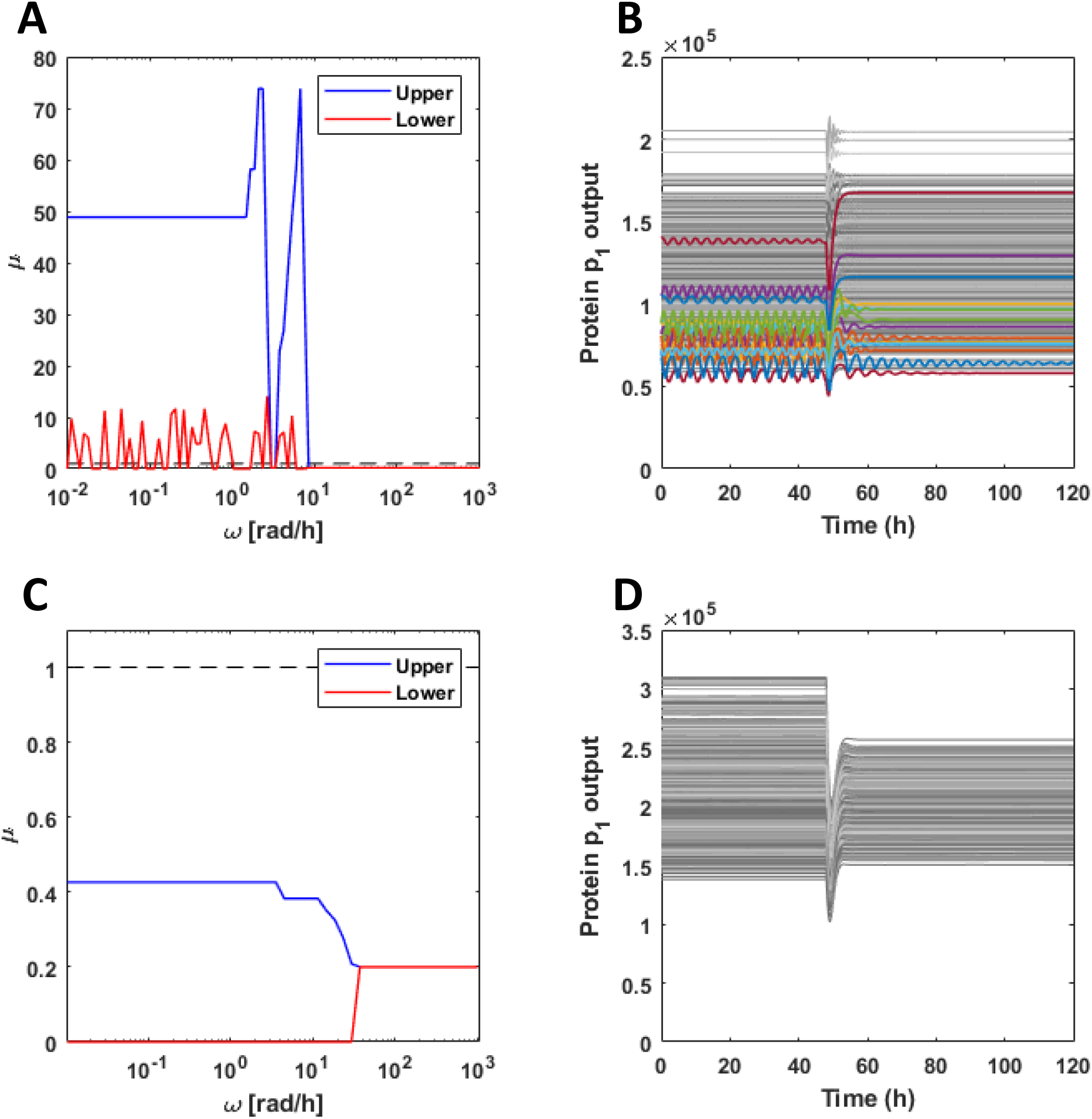
Robustness analysis of the dual control systems. The robustness of the two best control systems was quantified using an LFT-free *μ* analysis algorithm. The *μ*-analysis was carried out as described in the Methods. The estimates shown are scaled to show the value of *μ* for a maximum uncertainity of up to 20%. Monte carlo simulations of the non-linear models where carried out by simulating the action of perturbed controllers. The nominal controller designed as described in the main text was perturbed by random values up to 20%. The constitutive gene was initially simulated for a period before the second gene was induced (shown at 48 h). *N* = 1, 000 samplings. **(A)** μ estimate for the best decoupling design of UBER-OR. **(B)** Monte Carlo samplings show some controller parametrisations cause oscillations in process genes. These traces are highlighted in colour while stable results are shown in grey. N = 1, 000. **(C)** μ estimate for the best decoupling design of FRAG-OR. **(D)** Monte Carlo samplings do not identify any unstable controllers. *N* = 1,000.

In contrast, the best performing design of the FRAG-OR controller has high levels of robustness, with *μ* estimated to be between 0.20 and 0.43 for an uncertainty level of ±20% in all parameters (Figure 4C). Monte Carlo simulations of the full non-linear model at this level of uncertainty also failed to find a single unstable case, and show acceptable variations in performance (Figure 4D). Extending this analysis to the other designs for this controller architecture shows significantly greater levels of robustness in general; with only 16 of 100 controllers having an upper estimate of *μ* of more than 1 at 10% uncertainty (Figure S5D), and none of the controllers being guaranteed to be unstable for up to 10% uncertaintly (i.e. the lower estimate of *μ* is always less than one). This rises to only 4 controller designs with up to 20% uncertainty (Figure S5C).

## 3 Discussion

Competition for shared cellular resources results in the emergence of non-regulatory interactions between circuit genes. By exploiting recent advances in the creation of orthogonal biological components, RNA polymerase allocation controllers can be used to decouple co-expressed genes at the the transcriptional level and ribosomal allocation controllers can be used to decouple co-expressed genes at the translational level. Here, we investigated the hypothesis that by combining these control systems we could decouple genes at both the transcriptional and translational levels *simultaneously*. Interestingly, we find that simply combining separately designed (and functioning) transcriptional and translational controllers can result in an unstable or non-functional dual controller, e.g. implementing the universal bacterial expression resource transcriptional controller with the orthogonal ribosome controller produced sustained oscillations in circuit genes, while combining a translational controller with a fragment RNA polymerase controller abolishes the translational decoupling. From a Control Engineering viewpoint, this is not in fact surprising, as it is well known that combining several (separately stable) feedback loops into a single multivariable control system can readily produce instability or significant performance degradation due to interactions between the different controllers [25].

The standard approach used by control engineers to deal with this problem is to design and analyse all elements of the overall control system simultaneously using multivariable methods, [26], and this is the approach we adopted here, evaluating a large number of potential designs based on two alternative (biologically feasible) dual controller architectures. We find that UBER-OR, based on using the UBER system for transcriptional resource allocation, shows better (nominal) decoupling performance, whereas FRAG-OR, based on the fragmented RNA polymerase has higher levels of gene expression, indicating a tradeoff between these two performance objectives. The translational elements of both dual controllers show similar design principles to those identified in [18], with the best decoupling provided by higher copy numbers for the repressor *p_f_* gene, which itself should be multimeric and produced from strong ribosome binding sites. The RNA polymerase component of both controllers should be expressed at moderate levels and its regulator (*p_q_* in the case of the UBER-OR architecture and the *α*-fragment in the case of FRAG-OR) expressed with a weaker RBS to reduce competition for ribosomes.

The controllers were designed assuming that all system parameter values are precisely known. However, in reality biological parameters are rarely accurately known. Biological measurements are reported within possible error ranges and, even where these are small, introducing parts into new contexts, either DNA sequence or host genetic background, can cause unpredictable changes in component dynamics. These uncertainties can introduce instabilities such as oscillations in circuit genes. We utilised the *μ* analysis tool from Robust Control Theory to rigorously quantify the robustness of each design (both controller architecture and implementation) to parametric uncertainty. *μ* analysis offers several advantages over standard Monte Carlo sampling techniques, including a higher likelihood of finding the smallest destabilising level of uncertainty, and faster computation times (*μ* evaluates a design’s robustness to all potential uncertainty levels, while Monte Carlo sampling needs to be repeated at each different level of uncertainty).

Our analysis suggests that while controllers with UBER-OR architecture have the potential to deliver the highest levels of decoupling performance, they have inherently poor robustness (they are prone to become unstable if multiple parameters are not precisely tuned), thus motivating further research to explore potential improvements to this controller architecture. We note that the underlying universal bacterial expression resource shows some instability in its design and this may be due to the presence of a strong autocatalytic activator (the positive feedback of the orthogonal RNA polymerase) which produces its own inhibitor (the *p_q_* protein). This activation-inhibition motif is known to be prone to the emergence of oscillations, with designs that are known to be stable rapidly become unstable when parameters change slightly [27, 28]. In contrast, dual controllers based on the architecture employing a fragment RNA polymerase controller exhibited lower levels of nominal decoupling performance but much higher levels of robustness across multiple different potential implementations. Again, this tradeoff between nominal performance and robust stability is widely observed in engineered control systems [29], and thus may simply reflect the inherent limitations of biologically realistic schemes for dynamic resource allocation.

## 4 Methods

### 4.1 Circuit model with competition for host cellular resources

The core of the process model represents transcription and translation of a single unregulated gene and takes account of competition for cellular resources. Each circuit gene’s promoter, *g_i_*, is reversibly bound by the host’s RNA polymerase, *P_h_*, to produce the transcription complex, *x_i_*. This produces the mRNA, *m_i_*, at rate *τ_i_*. Completion of transcription also liberates the free promoter and RNA polymerase:

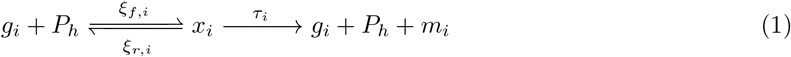

The mRNA reversibly binds to host ribosomes, *R_h_*, to produce the intermediate translational complex, *c_i_*. This produces protein, *p_i_*, at rate *γ_i_*. Completion of translation also liberates the mRNA and ribosome:

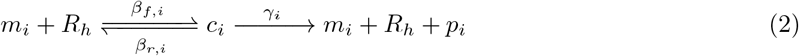

We assume that all species dilute due to growth while mRNAs also decay [30]:

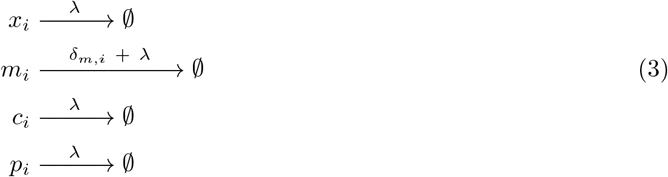

By applying the Law of Mass Action we can derive the following dynamics:

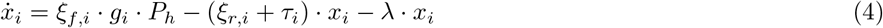

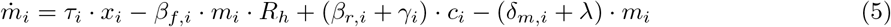

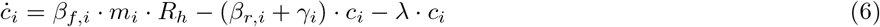

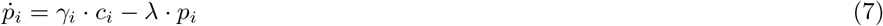

Due to their own control mechanisms the copy number of each plasmid and therefore gene *g_i_* remains constant such that a conservation law can be applied. The total number of promoters for each gene is constant *ω_i_* = *g_i_* + *x_i_*. From this we calculate the concentration of the free promoter:

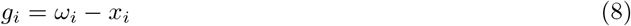

Similarly, the host’s internal control mechanisms maintain the total number of RNA polymerases and ribosomes for any given growth rate and as such we can calculate the concentration of free resources:

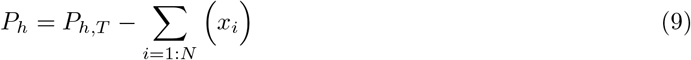

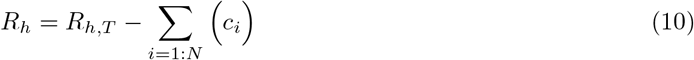

where *P_h,T_* and *R_h,T_* are the total concentrations of the host’s RNA polymerase and ribosomes respectively.

Induction of each circuit module can be simulated by varying the concentration of each promoter *ω_i_*. Note that this allows us to neglect the regulation of each circuit promoter and so simplify the model.

### 4.2 Resource allocation controller models

#### 4.2.1 Kushwaha and Salis transcriptional resource allocation controller model

The universal bacterial expression resource developed by Kushwaha and Salis utilises an orthogonal RNA ploymerase, *P_o_*, as the new circuit specific transcriptional resource. The RNA polymerase transcribes itself and a second repressor protein, *p_Q_*, which inhibits the o-RNAP promoter, *g_p_*, to produce a sequestered complex *k_p_*:

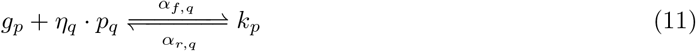

This species dilutes due to growth as in [30]:

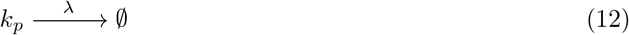

We assume that the production of the orthogonal RNA polymerase is leaky:

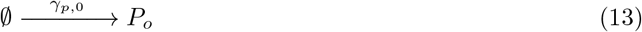

The transcription and translation of *P_o_* and *p_q_* mirror the chemical reaction networks in Section ??. Applying the Law of Mass Action yields the following dynamics for the production of the o-RNAP:

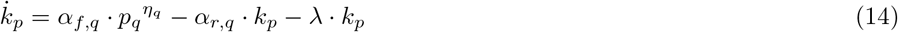

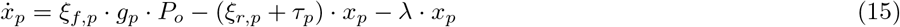

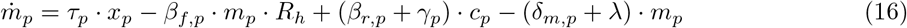

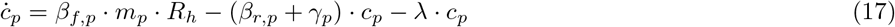

The dynamics of the orthogonal RNA polymerase which transcribes circuit genes are given by:

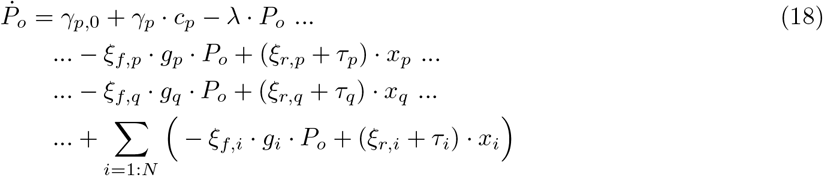

The production of the controller protein, *p_q_*, follows the dynamics for circuit proteins in (??) to (6) except that the host RNAP, *P_h_*, is replaced with the o-RNAP, *P_o_*. The translational machinery remains the host ribosome, *R_h_*:

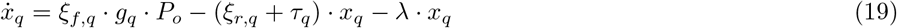

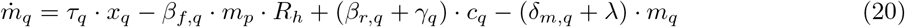

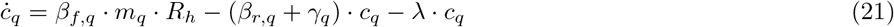

The dynamics of the final protein, including the *g_p_*-*p_q_* repression interaction, are given by:

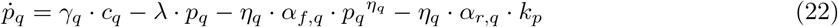

The concentration of free promoters of the orthogonal RNA polymerase and its regulator *p_q_* can be calculated from their respective total *ω_p_* and *ω_q_*:

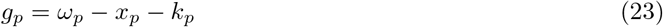

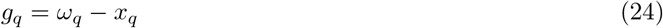

This controller utilises its own RNA polymerase for its own expression and for circuit gene transcription, therefore the host RNA polymerase is not utilised:

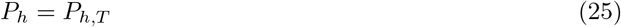

The controller utilises the host translation system for its own expression and expression of circuit genes:

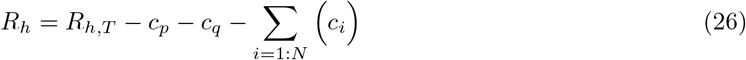

#### 4.2.2 The Segall-Shapiro *et al.* transcriptional resource allocation controller model

Segall-Shapiro *et al.* utilise a RNA polymerase which, rather than being made of up a single protein, is split into two components. The core *β* fragment (*p_c_*) is constitutively expressed and sets a ‘transcriptional budget’ which is then targeted by the expression of a ‘synthetic’ *σ* factor, *p_a_*. These two components bind to create the functional circuit-specific orthogonal RNA polymerase:

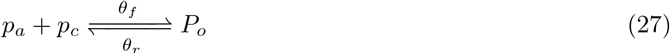

Again, we assume that the production of the orthogonal RNA polymerase is leaky:

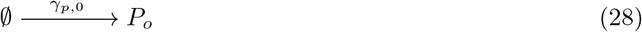

The production of each protein component follows the same dynamics as for circuit proteins. Applying the Law of Mass Action, including the production of RNAP in Eq. 27, gives:

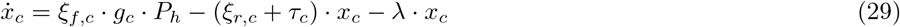

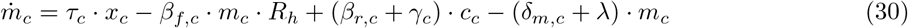

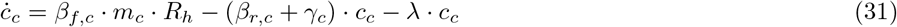

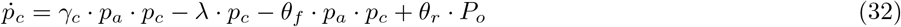

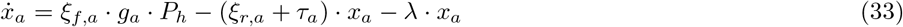

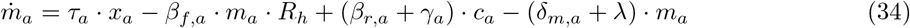

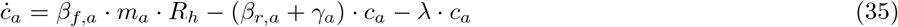

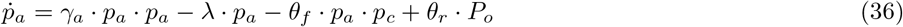

The dynamics of the orthogonal RNA polymerase which transcribes circuit genes are given by:

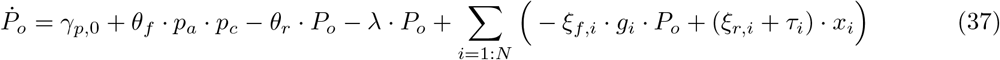

The concentration of free promoters of the core RNA polymerase and its targeting fragment can be calculated from their repsective total *ω_c_* and *ω_α_*:

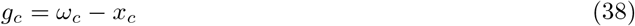

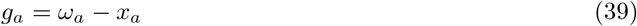

As described in [16], the promoters of the targeting fragment of the orthogonal split RNA polymerase are carried on the circuit plasmids and therefore their total copy number is the sum of the circuit promoters:

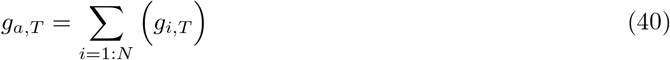

This controller utilises its own RNA polymerase for circuit gene expression but utilises the host’s for its own production such that the free host RNA polymerase is given by:

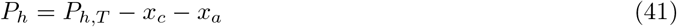

The controller utilises the host translation system for its own expression and expression of circuit genes:

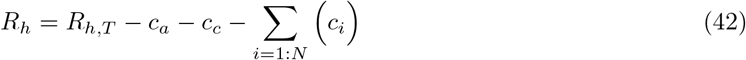

#### 4.2.3 Translational resource allocation controller model

We model the conversion of ribosomes between the host and orthogonal ribosome pool by considering the one step reaction between an orthogonal rRNA, *r*, and the host ribosome, *R_h_*, to produce orthogonal ribosomes, *R_o_*:

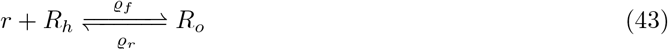

The rRNA and orthogonal ribosome are also subject to dilution:

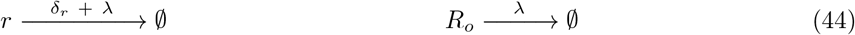

In [17], we developed a translational controller which dynamically allocates the distribution between host and orthogonal ribosomes by placing the o-rRNA gene, *g_r_* under the control of a protein, *p_f_*, which itself is translated by the orthogonal ribosome pool. The controller protein, *p_f_*, sequesters free rRNA promoters to an inactive complex, *k_r_*:

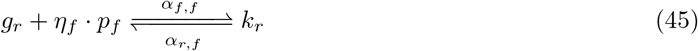

As with all other species, this dilutes due to growth as in [30]:

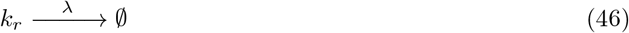

Applying the Law of Mass Action yields the following dynamics:

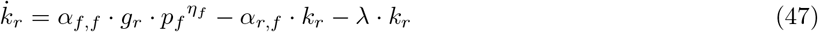

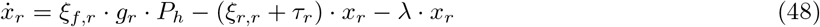

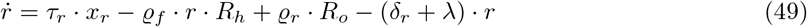

The dynamics of the orthogonal ribosome pool are given by:

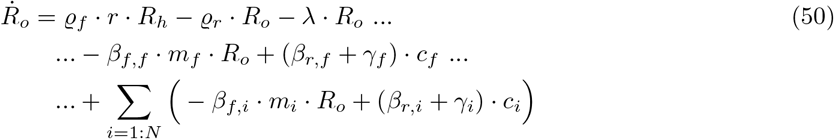

The production of the controller protein, *p_f_*, follows the same dynamics of circuit proteins, except the host ribosome, *R_h_*, is replaced with its orthogonal counterpart, *R_o_*:

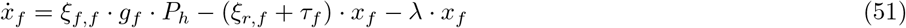

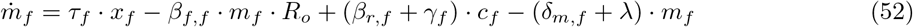

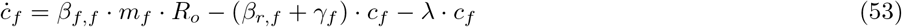

The dynamics of the inhibitory protein are:

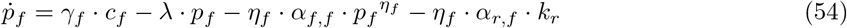

The concentration of free promoters of the o-rRNA and controller protein can be calculated from their respective total *ω_r_* and *ω_f_*:

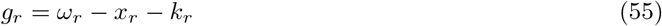

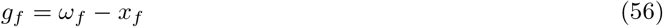

This controller utilises the host RNA polymerase for all transcription:

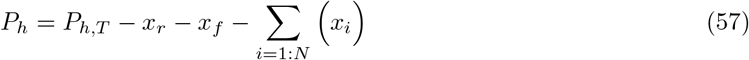

As described above the controller co-opts ribosomes from the host such that the free host ribosome pool is given by:

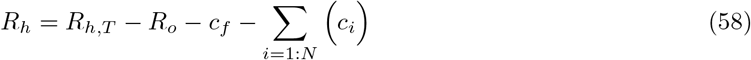

### 4.3 Dual transcriptional-translational controller models

We combine the two transcriptional controller with the translational controller as follows. In each case the host cellular resources in circuit (Eq. 4-7) are replaced with their orthogonal counterparts (*P_h_* ⟹ *P_o_* and *R_h_* ⟹ *R_o_* respectively). We also update the equations describing the use of host resources.

Combining the Kushwaha controller with the translational controller results in the following host resource usage:

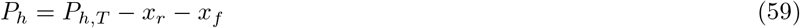

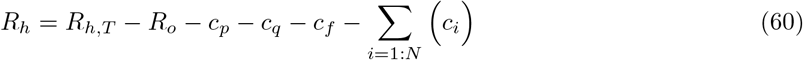

The designable parameters of this controller are: *ω_p_, ξ_f,p_, β_f,p_, ω_r_, ξ_f,r_, ω_f_, ξ_f,q_, β_f,q_, ω_q_, ξ_f,f_, β_f,f_, α_r,q_, η_q_, α_r,f_, η_f_*.

Combining the Segall-Shapiro controller with the translational controller results in the following host resource usage:

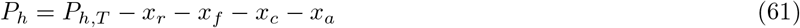

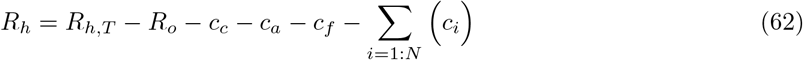

The designable parameters of this controller are: *ω_c_, ξ_f,c_, β_f,c_, ξ_f,α_, β_f,a_, ω_r_, ξ_f,r_, ω_f_, ξ_f,f_, β_f,f_, α_r,f_, η_f_*.

The equations needed to simulate the full models of each controller are shown in Table 1.

### 4.4 Designing experimentally implementable parameters

As discussed in [18], we set the total concentration of host RNA polymerase and ribosomes to 250 nM and 2,500 nM respectively assuming a constant growth rate of 1 h^-1^. The model assumes that each gene is bound by only one RNA polymerase and each mRNA is bound by one ribosome. However, *in vivo* each gene is transcribed by multiple RNA polymerase and each mRNA is translated by multiple ribosomes. We account for this in our model by increaseing the copy number of each gene and increaseing the mRNA production rate such that RNA polymerase and ribosome are subject to the the appropriate level of competition. From [18], take conservative estimates and increase gene copy numbers by 10 and the mRNA production rates by 20 throughout.

The strengths of promoters, RBS sites and protein DNA binding are often quoted as dissociation constants. Following from the definitions in [8, 11, 18], the RNAP-promoter, ribosome-RBS and transcription factor dissociation constants are given by *k_X_, k_L_* and *μ* respectively:

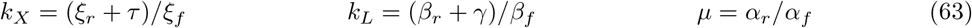

As our model contains resource turnover (as a specific consequence of modelling control of resources) we do not make the quasi-steady state assumption as in [8, 11, 18] and so the individual binding/unbinding rate constants remain in their original form in the ODEs. These lumped dissociation constants do not appear. Therefore, in order to vary *k_X_* and *k_L_* we vary *ξ_f_* and *β_f_*. From [18], *k_X_* ranges from 5 nM to 1,000 nM and assuming that *β_r_* + *γ* ≈ 1100 then *ξ_f_* varies from 11 and 2200. Similarly, *k_L_* ranges from 10^4^ to 10^7^ nM with *β_r_* + *γ* ≈ 10^6^ then *β_f_* varies from 0.1 to 100.

We assume that promoters can be carried on a low (10 nM), medium (100 nM) or high (500 nM) copy number plasmids.

For simplicity we limit the choice of repressors for *p_f_* and *p_q_* to the commonly used repressors lacI, cI from bacteriophage λ and tetR. This limits the values of the promoter-RNA polymerase dissocation constant, transcription factor dissociation constant and multimerisation. Note that we assume that the transcription factor binding rate (*α_f_*) is 1 and therefore *α_r_* is set to the value of the dissociation constant, *μ*. The values of these parameters are listed in Table 2. All other parameters are shown in Table 3.

**Table 2:** Transcription factor parameters. Note that when expressed using the orthogonal RNA polymerase (such as in the universal bacterial expression resource) we set *k_X_* = 200 nM and therefore set *ξ_f_* to 55 (nM · h)^-1^

| Name | $k_X$ nM | $\xi_f$ (nM · h)<sup>−1</sup> | $\alpha_r$ ($\approx \mu$) (nM) | $\eta_f$ |
| --- | --- | --- | --- | --- |
| lacI | 550 | 20.0 | 0.02 | 4 |
| tetR | 350 | 31.4 | 167 | 1 |
| cI | 100 | 110.0 | 22 | 2 |

**Table 3:**
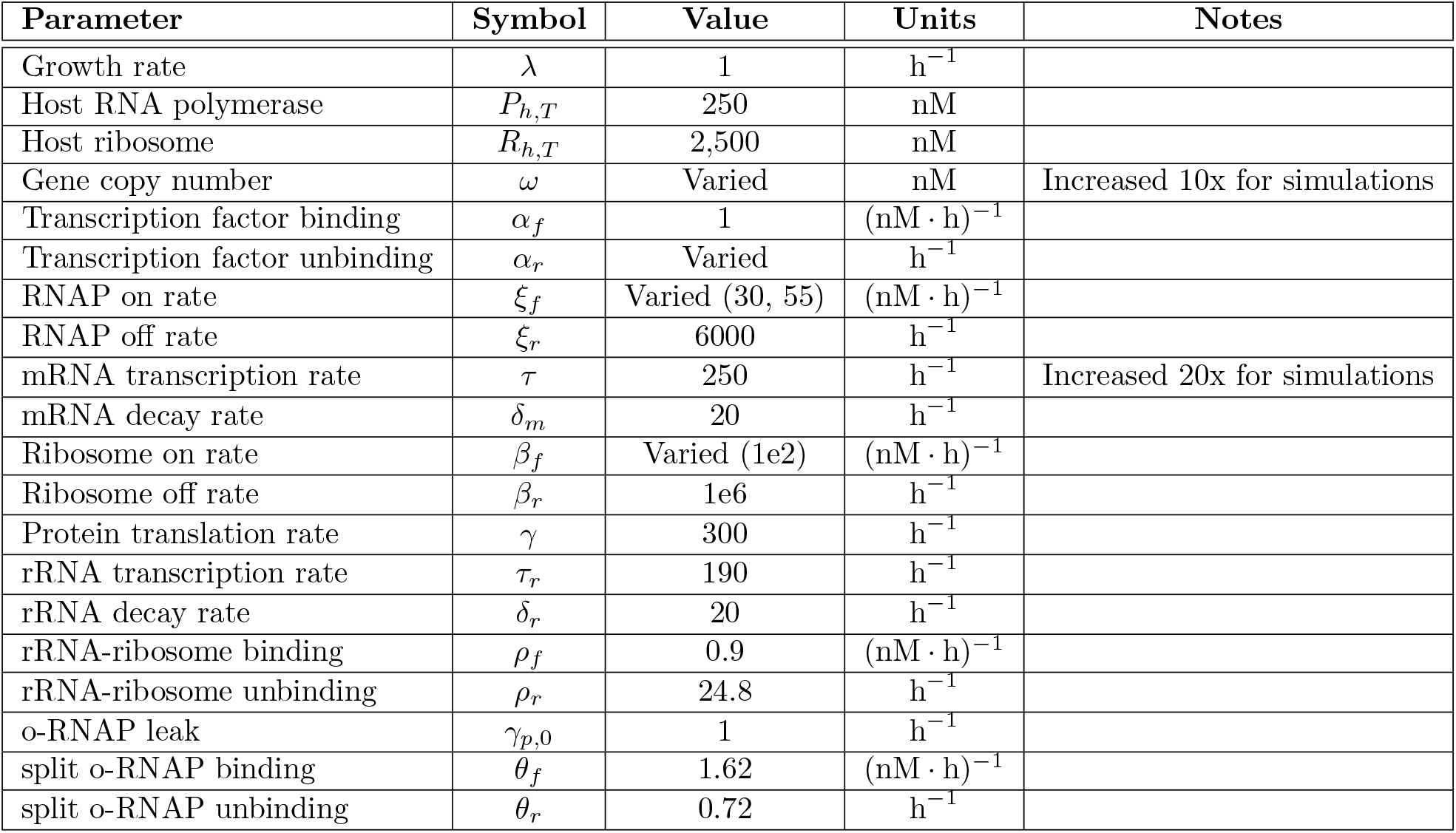
Parameters common to all models. Note that when genes are expressed using the orthogonal RNA polymerase (such as in the universal bacterial expression resource) we set *k_X_* = 200 nM and therefore set f to 55 (nM · h)^-1^. When genes are expressed using the host RNA polymerase we set *k_X_* = 366 nM and thefore set *ξ_f_* to 30 (nM · h)^-1^.

| Parameter | Symbol | Value | Units | Notes |
| --- | --- | --- | --- | --- |
| Growth rate | $\lambda$ | 1 | $\text{h}^{-1}$ | |
| Host RNA polymerase | $P_{h,T}$ | 250 | nM | |
| Host ribosome | $R_{h,T}$ | 2,500 | nM | |
| Gene copy number | $\omega$ | Varied | nM | Increased 10x for simulations |
| Transcription factor binding | $\alpha_f$ | 1 | $(\text{nM} \cdot \text{h})^{-1}$ | |
| Transcription factor unbinding | $\alpha_r$ | Varied | $\text{h}^{-1}$ | |
| RNAP on rate | $\xi_f$ | Varied (30, 55) | $(\text{nM} \cdot \text{h})^{-1}$ | |
| RNAP off rate | $\xi_r$ | 6000 | $\text{h}^{-1}$ | |
| mRNA transcription rate | $\tau$ | 250 | $\text{h}^{-1}$ | Increased 20x for simulations |
| mRNA decay rate | $\delta_m$ | 20 | $\text{h}^{-1}$ | |
| Ribosome on rate | $\beta_f$ | Varied (1e2) | $(\text{nM} \cdot \text{h})^{-1}$ | |
| Ribosome off rate | $\beta_r$ | 1e6 | $\text{h}^{-1}$ | |
| Protein translation rate | $\gamma$ | 300 | $\text{h}^{-1}$ | |
| rRNA transcription rate | $\tau_r$ | 190 | $\text{h}^{-1}$ | |
| rRNA decay rate | $\delta_r$ | 20 | $\text{h}^{-1}$ | |
| rRNA-ribosome binding | $\rho_f$ | 0.9 | $(\text{nM} \cdot \text{h})^{-1}$ | |
| rRNA-ribosome unbinding | $\rho_r$ | 24.8 | $\text{h}^{-1}$ | |
| o-RNAP leak | $\gamma_{p,0}$ | 1 | $\text{h}^{-1}$ | |
| split o-RNAP binding | $\theta_f$ | 1.62 | $(\text{nM} \cdot \text{h})^{-1}$ | |
| split o-RNAP unbinding | $\theta_r$ | 0.72 | $\text{h}^{-1}$ | |

### 4.5 Controller design and other numerical methods

We developed a range of experimentally feasible designs based on biologically realisable promoter and RBS strengths, gene copy numbers, and the common repressors tetR, lacI and cI (as discussed above). We solved these ODEs for each controller numerically using the inbuilt ordinary differential equation solver ode23s in MATLAB 2019a. We assess the performance of these controllers by simulating the response to two sequential large step inputs (i.e. simulating the induction of genes sequentially from a high copy number plasmid). To determine stability (or otherwise) of the controllers, we numerically calculate the jacobian and eigvenvalues in each part of the simulation (i.e. before induction, after induction of the first gene and after the induction of the second gene) and remove controllers which have positive real eigenvalues. We also remove controllers which inhibit the process output sufficiently to reduce process output to near zero.

To assess the performance of each controller, we calculate coupling at both the transcriptional and translational levels by assessing the fall in the first circuit gene (mRNA *m*_1_ and protein *p*_1_ respectively) in response to the induction of the second circuit gene by comparing the values at the time of the second gene induction *t_ind_* and the end of the simulate *t_max_*:

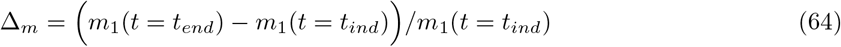

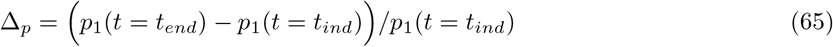

We scale Δ_*m*_ and Δ_*p*_ by the maximum value obtained. To create a single performance metric we consider the behaviour of the controllers in a two dimensional plane whose axes are the scaled values of Δ_*m*_ and Δ_*p*_. The origin (0,0) represents perfect decoupling at both the transcriptional and translational levels. We calculate the Euclidean distance of each point from the origin:

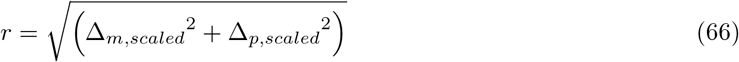

We sort the controller by *r* and take the top *N* = 100 controllers which represent those which show the best decoupling performance. We call this metric the 2D score. We assess these controllers further in terms of both decoupling and protein output.

### 4.6 LFT-free *μ* analysis

We define the following two vectors containing the variables and dynamics respectively **y** and **ẏ** where both are *n* × 1 vectors containing the species variables and dynamics as required for each system. We define **ȳ** to be the solution to **ẏ** = **0**. Given the nonlinearity of **ẏ** we find **y** by solving **ẏ** numerically as described above. We define a small perturbation around **ȳ** as **x** = **y** – **ȳ** and linearise the model around this point:

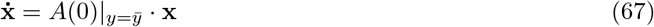

where *A*(0) is the Jacobian of the certain (‘nominal’) system. The uncertain Jacobian is *A*(*δ*) where *δ* is a vector containing the *n* × 1 uncertainties. We represent the uncertain system as:

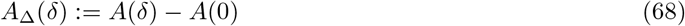

*A*_Δ_(*δ*) can be calculated numerically by sampling 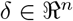. The linearisation can now be represented as:

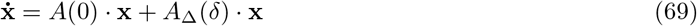

As *μ* analysis tools have been developed based on the transfer function in the frequency domain we take the Laplace transform to represent **x**(*t*) in the frequency domain *X*(*s*):

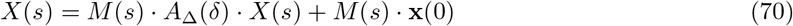

where *M*(*s*) = [*s* · *I* – *A*(0)]^−1^ and **x**(0) is the initial condition of **x**(*t*).

The robustness problem is now formulated as a search for the smallest value of *δ* which satisfies [21, 22]:

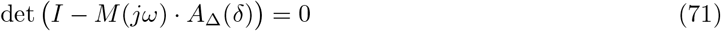

for all sampled frequencies *ω* ∈ [0,inf]. This corresponds as the intersection between the functions *f_R_* and *f_I_*:

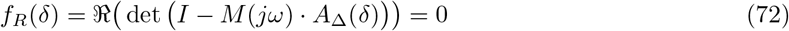

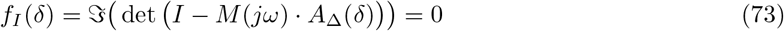

where 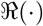 and 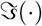 are the real and imaginary components of the complex number.

By definition for the nominal system *A*_Δ_(0) = 0, in this case *I* – *M*(*jω*) · *A*_Δ_(0) = *I*, i.e. the identify matrix, and the determinant is 1. Therefore the *f_R_*(0) = 1 and *f_I_*(0) = 0. Therefore, the in uncertainty space *f_I_* must always pass though the origin and *f_R_* must not. The uncertainty space can then be divided into four sections (Figure S6):

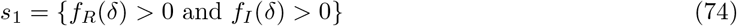

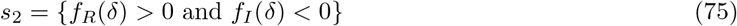

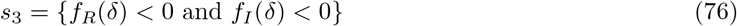

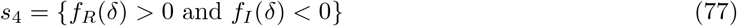

This intersection point can be identified efficiently using the algorithms described in [24]. The algorithm functions by considering two boxes in uncertainty space - one which must contain all four s points (this gives the lower bound – i.e. *μ* must be within the box) and one which much contain only three of the s points (this gives the upper bound – i.e. *μ* cannot be within this box) (Figure S6). The inverse of the size of each box gives the respective *μ* bound. The algorithm was run with the following settings: the number of points of each box: 5,000; tolerance: 0.0001; frequencies: 36 evenly spaced (log scale) points from 10^-2^ to 10^3^.

## 5 Author contributions

Conceptualization, Methodology, Formal analysis and Writing (original draft): APSD. Supervision, Funding acquisition: DGB. Writing (review and editing): APSD and DGB.

## 6 Acknowledgements

APSD is grateful to Jongrae Kim of the University of Leeds for initial discussions around the instability observed in the UBER-OR controller and for providing the MATLAB implementation of the LFT-free μ-analysis algorithms. APSD and DGB acknowledges funding from The Leverhulme Trust (grant RPG-2017-284) and the high performance computing facilities provided by Warwick Integrative Synthetic Biology Centre (grant BB/M017982/1).

## References

[1] J. A. Brophy and C. A. Voigt, “Principles of genetic circuit design.” Nature Methods, vol. 11, no. 5, pp. 508–20, 2014.

[2] S. Cardinale and A. P. Arkin, “Contextualizing context for synthetic biology - identifying causes of failure of synthetic biological systems,” Biotechnology Journal, vol. 7, no. 7, pp. 856–866, 2012.

[3] O. Borkowski, F. Ceroni, G.-b. Stan, and T. Ellis, “Overloaded and stressed: whole-cell considerations for bacterial synthetic biology,” Current Opinion in Microbiology, vol. 33, pp. 123–130, 2016.

[4] F. Moser, N. J. Broers, S. Hartmans, A. Tamsir, R. Kerkman, J. A. Roubos, R. Bovenberg, and C. A. Voigt, “Genetic circuit performance under conditions relevant for industrial bioreactors,” ACS Synthetic Biology, vol. 1, no. 11, pp. 555–564, 2012.

[5] C. Vilanova, K. Tanner, P. Dorado-Morales, P. Villaescusa, D. Chugani, A. Frías, E. Segredo, X. Molero, M. Fritschi, L. Morales, D. Ramón, C. Peña, J. Peretó, and M. Porcar, “Standards not that standard.” Journal of biological engineering, vol. 9, e17, 2015.

[6] M. Kushwaha and H. M. Salis, “A portable expression resource for engineering cross-species genetic circuits and pathways,” Nature Communications, vol. 6, e7832, 2015.

[7] F. Ceroni, R. Algar, G.-B. Stan, and T. Ellis, “Quantifying cellular capacity identifies gene expression designs with reduced burden,” Nature Methods, vol. 12, no. 5, pp. 415–423, 2015.

[8] A. Gyorgy, J. I. Jiménez, J. Yazbek, H.-H. Huang, H. Chung, R. Weiss, and D. Del Vecchio, “Isocost Lines Describe the Cellular Economy of Genetic Circuits,” Biophysical Journal, vol. 109, no. 3, pp. 639–646, 2015.

[9] M. Carbonell-Ballestero, E. Garcia-Ramallo, R. Montanez, C. Rodriguez-Caso, and J. Marcia, “Dealing with the genetic load in bacterial synthetic biology circuits: convergences with the Ohm’s law,” Nucleic Acids Research, vol. 44, no. 1 pp. 496–507, 2015.

[10] T. E. Gorochowski, I. Avcilar-Kucukgoze, R. A. Bovenberg, J. A. Roubos, and Z. Ignatova, “A minimal model of ribosome allocation dynamics captures trade-offs in expression between endogenous and synthetic genes,” ACS Synthetic Biology, vol. 5, no. 7, pp. 710–720, 2016.

[11] Y. Qian, H.-H. Huang, J. Jiménez, and D. Del Vecchio, “Resource competition shapes the response of genetic circuits,” ACS Synthetic Biology, vol. 6, no. 7, pp. 1263–1272, 2017.

[12] R. D. Jones, Y. Qian, B. Diandreth, V. Siciliano, and J. Huh, “An endoribonuclease-based incoherent feedforward loop for decoupling resource-limited genetic modules,” bioXriv pre-print doi:10.1101/867028, 2019.

[13] T. Frei, F. Cella, F. Tedeschi, J. Gutierrez, G.-B. V. Stan, M. H. Khammash, and V. Siciliano, “Characterization, modelling and mitigation of gene expression burden in mammalian cells,” bioXriv pre-print doi:10.1101/867549, 2019.

[14] W. An and J. W. Chin, “Synthesis of orthogonal transcription- translation networks,” Proceedings of the National Academy of Sciences, vol. 106, no. 21, pp. 8477–8482, 2009.

[15] C. C. Liu, M. C. Jewett, J. W. Chin, and C. A. Voigt, “Towards an orthogonal central dogma,” Nature Chemical Biology, vol. 14, no. 2, pp. 103–106, 2018.

[16] T. H. Segall-Shapiro, A. J. Meyer, A. D. Ellington, E. D. Sontag, and C. A. Voigt, “A ‘resource allocator’ for transcription based on a highly fragmented T7 RNA polymerase.” Molecular Systems Biology, vol. 10, no. 7, p. 742, 2014. [Online]. Available: http://www.ncbi.nlm.nih.gov/pubmed/25080493

[17] A. P. S. Darlington, J. Kim, J. I. Jiménez, and D. G. Bates, “Dynamic allocation of orthogonal ribosomes facilitates uncoupling of co-expressed genes,” Nature Communications, vol. 9, e695, 2018.

[18] A. P. S. Darlington, J. Kim, J. I. Jiménez, and D. G. Bates, “Engineering translational resource allocation controllers: Mechanistic models, design guidelines, and potential biological implementations,” ACS Synthetic Biology, vol. 7, no. 11, pp. 2485–2496, 2018.

[19] A. Tsigkinopoulou, S. M. Baker, and R. Breitling, “Respectful modeling: Addressing uncertainty in dynamic system models for molecular biology,” Trends in Biotechnology, vol. 35, no. 6, pp. 518–529, 2017.

[20] C. Zhang, R. Tsoi, and L. You, “Addressing biological uncertainties in engineering gene circuits.” Integrative biology, vol. 8, no. 4, pp. 456–564, 2015.

[21] J. Doyle, “Analysis of Feedback Systems With Structured Uncertainties.” IEE Proceedings D: Control Theory and Applications, vol. 129, no. 6, pp. 242–250, 1982.

[22] J. Doyle, A. Packard, and K. Zhou, “Review of LFTs, LMIs and μ,” in Proceedings of the 30th IEEE Conference on Decision and Control, Brighton, UK, pp. 1227–1232, 1991.

[23] Y.-b. Zhao, J. Kim, and D. G. Bates, “LFT-free μ-analysis of LTI / LPTV systems,” in IEEE International Symposium on Computer-Aided Control System Design, Denver, CO, USA, pp. 638–643, 2011.

[24] A. P. S. Darlington, J. Kim, and D. G. Bates, “Robustness analysis of a synthetic translational resource allocation controller”, IEEE Control Systems Letters, vol. 3, no. 2, pp. 266–271, 2019.

[25] D. Bates and I. Postlethwaite, Robust Multivariable Control of Aerospace S’ystems, Ios Pr Inc., 2002.

[26] C. Cosentino and D. Bates, Feedback Control in Systems Biology, CRC Press, 2011.

[27] M. R. Atkinson, M. A. Savageau, J. T. Myers and A. J. Ninfa, “Development of genetic circuitry exhibiting toggle switch or oscillatory behavior in Escherichia coli”, Cell, vol. 113, no. 5, pp. 597–607, 2003.

[28] S. Jayanthi and D. del Vecchio, “Tuning genetic clocks employing DNA binding sites”, PLOS ONE, vol. 7, no. 7, e41019, 2012.

[29] S. Skogestad and I. Postlethwaite, Multivariable feedback control - Analysis and design, 2nd ed., Wiley, 2005.

[30] E. J. Hancock, G.-B. V. Stan, J. A. J. Arpino, and A. Papachristodoulou, “Simplified mechanistic models of gene regulation for analysis and design,” Journal of The Royal Society Interface, vol. 12, e20150312, 2015.

