## Supplementary material for "Design trade-offs and robust architectures for combined transcriptional and translational resource allocation controllers"

February 2020

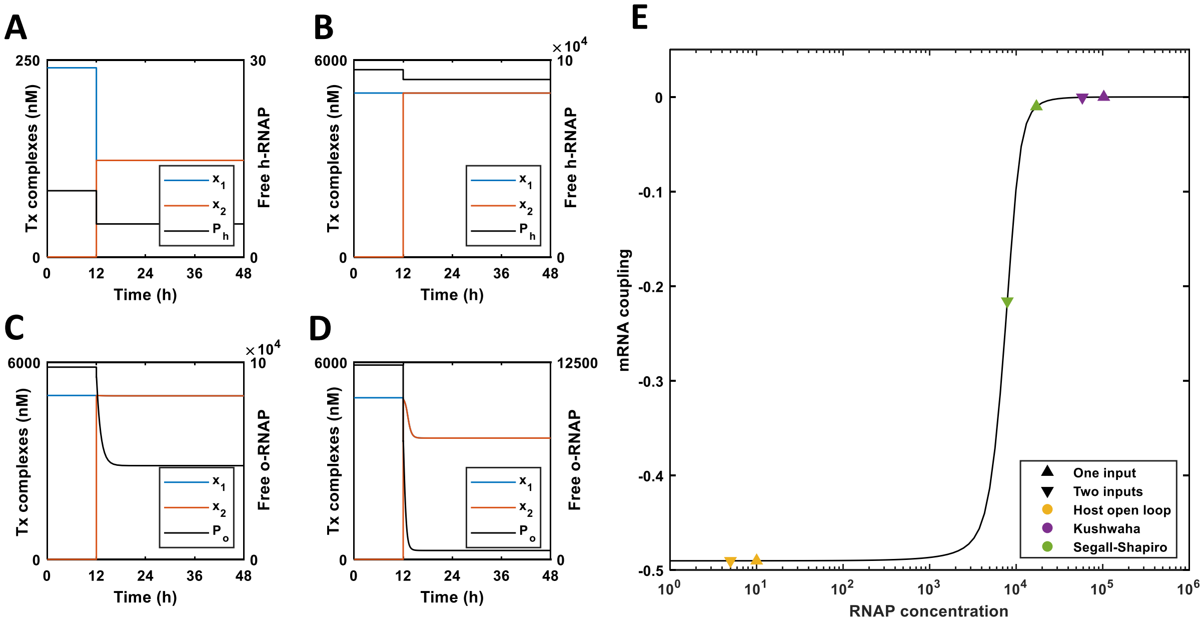


Figure S1: **Decoupling mechanism of the transcriptional controllers.** Simulations showing the ability of the transcriptional controllers to reject disturbances at the transcriptional level. **(A)** Host RNA polymerase at the biologically feasible concentration. **(B)** Increasing the host RNA polymerase allows decoupling in the absence of control but this is not through a biologically feasible mechanism. **(C)** Decoupling though the universal bacterial expression resource. **(D)** Decoupling though the fragmented RNA polymerase resource allocator. **(E)** Increasing RNA polymerase concentration in the absence of any control mechanism results in reduced mRNA coupling where $m_1$ is maintained while $m_2$ is induced.

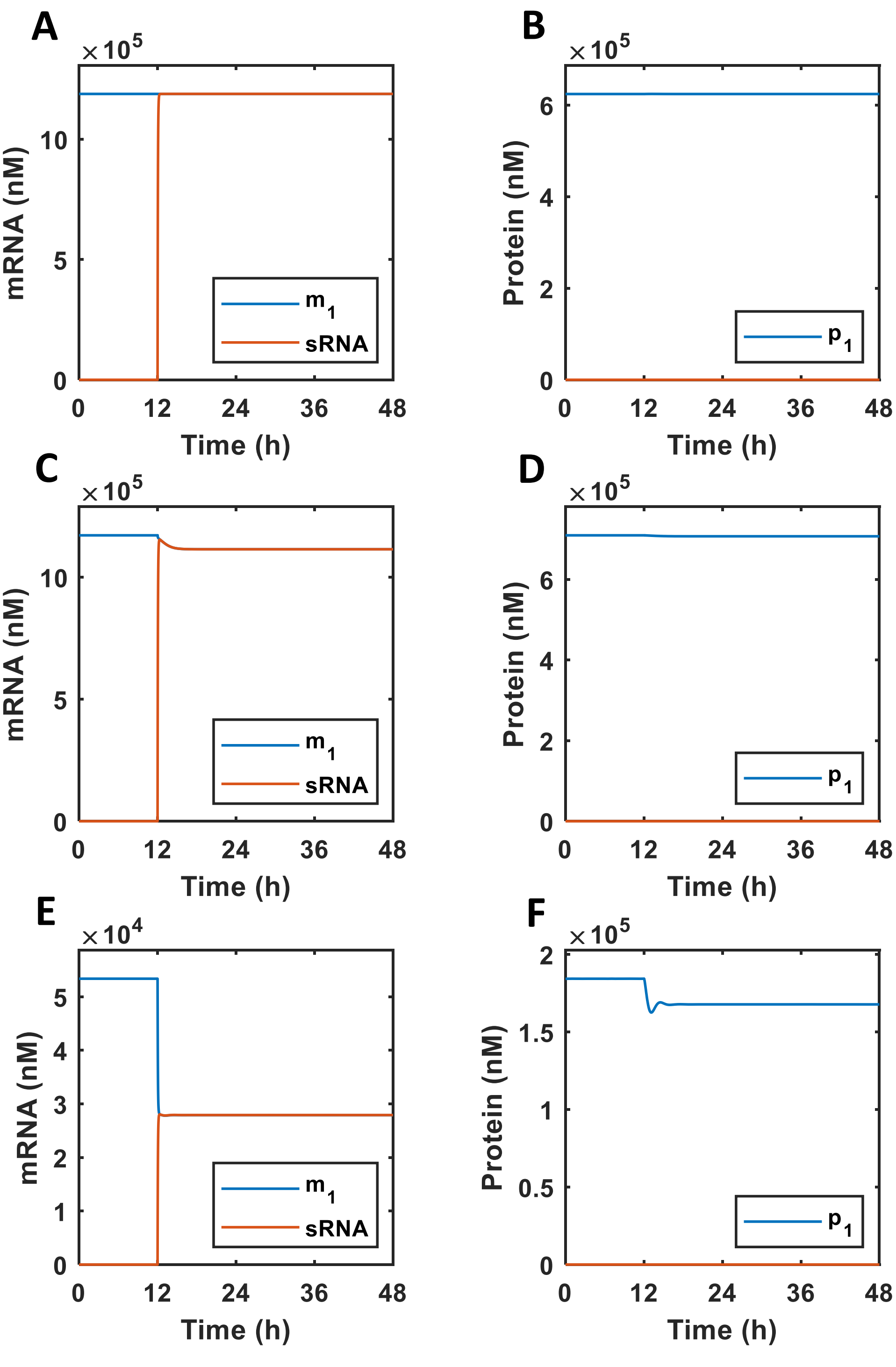


Figure S2: **Gene decoupling by transcriptional and translational controllers in response to a transcriptional disturbance.** The behaviour of candidate transcriptional and translational control systems in isolation. We simulated the ability of each prototype control system to decouple co-expressed genes; one encoding a protein $p_1$ and the other an sRNA. We consider the impact on a constitutively expressed gene ($g_1$) of the induction of a second gene ($g_2$) at 12 h. This results in a transcriptional only disturbance (as promoters compete for RNA polymerase). **(A)** The UBER controller successfully mitigates the transcriptional disturbance applied at 12 h. **(C)** There is no change in the level of $p_1$. **(D)** The fragmented RNA polymerase controller is able to mitigate the transcriptional disturbance at 12 h to some extent with the fall in $m_1$ only being 15% rather than 50%. **(D)** There is a negligible impact on the protein $p_1$. **(E)** The translational controller has no impact on the transcriptional disturbance with $m_1$ falling by 50% upon activation of the second protein-encoding gene. **(F)** Coupling remains at the protein level due to competition for RNA polymereases.

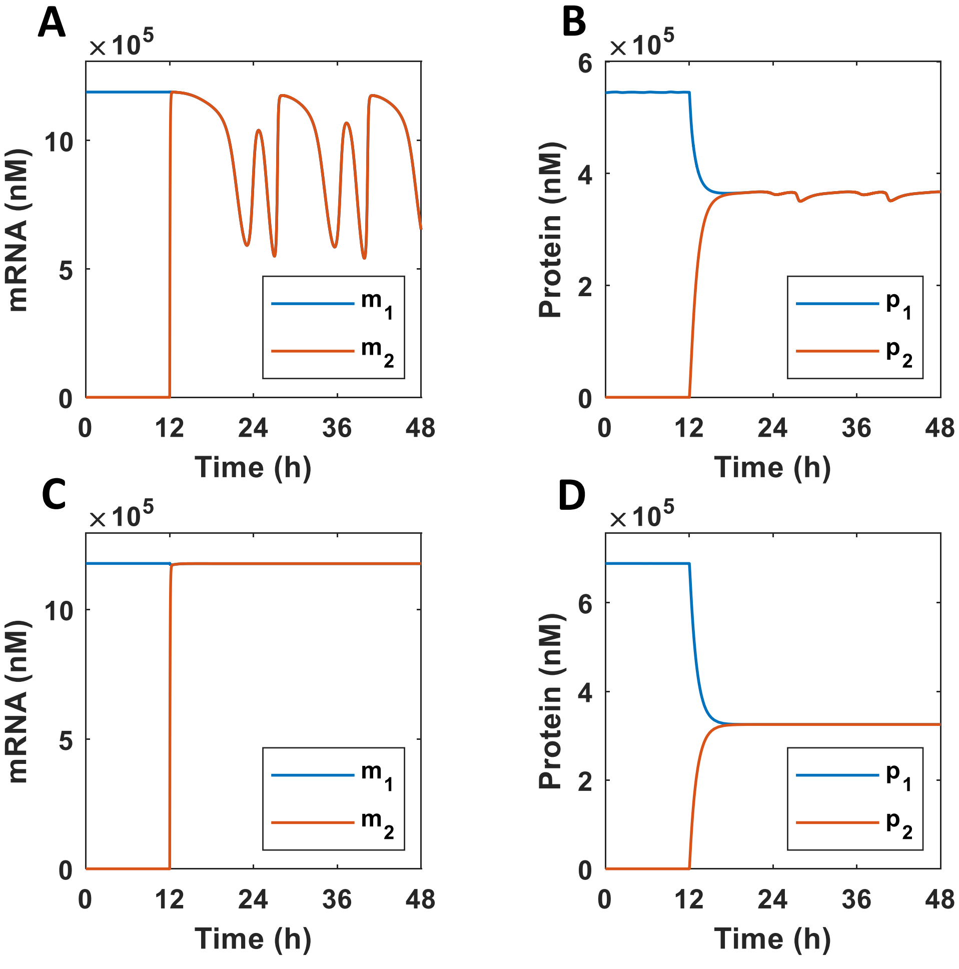


Figure S3: **Combining the transcriptional and translational controllers into new dual control systems.** The behaviour of the controllers depicted in Figure 1 when combined into dual control systems (as depicted in Figure 2). **(A)** The combined control system based on the universal bacterial expression resource and the translational controller (UBER-OR) results in significant oscillations at the transcriptional level. **(B)** The oscillations of the mRNA in (A) propagate to the translational level. **(C)** The combined control system based on the fragmented RNA polymerase and the translational controller (FRAG-OR) results in decoupling at the transcriptional level. **(D)** The combined control system does not impact protein-level coupling.

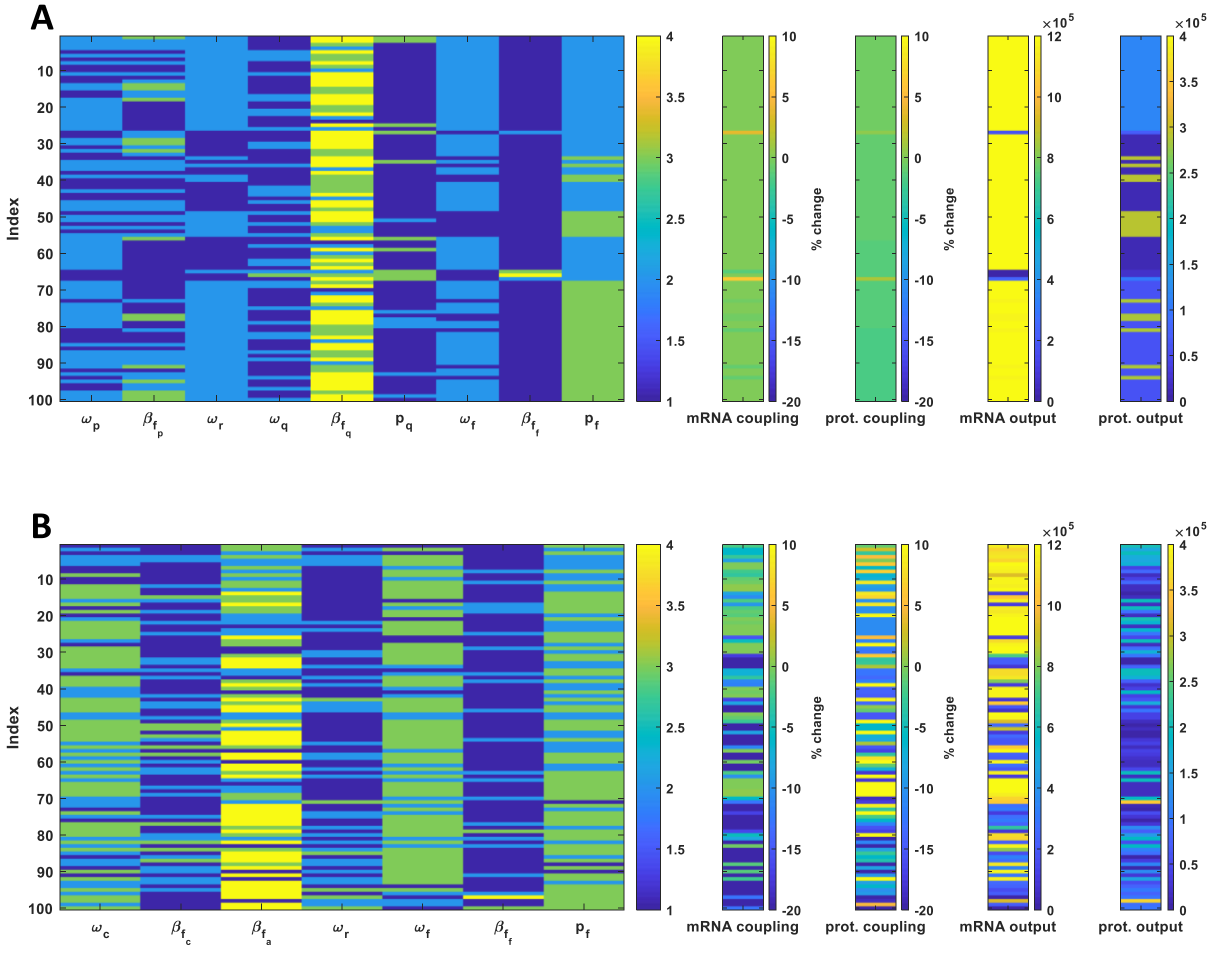


Figure S4: **Designs and performance of the best 100 controllers of each architecture.** The best parameteric designs of each controller where determined as outlined in the main text. As described the main text this design space is discrete rather than continuous. The parameter value colour corresponds to the index in the design vector. $\omega$ parameters can take the value 10, 100 or 500 nM. $\beta_f$ can take the values from the vector [100 10 1 0.1] which corresponds to a ribosome dissociation constant vector of $[10^4\ 10^5\ 10^6\ 10^7]$ nM. The proteins $p_q$ or $p_f$ can either be tetR (1), lacI (2) or cI (3) and this selection sets the values of the target RNA polymerase–promoter on rate $\xi_f$, the protein–gene off rate $\alpha_r$ and the co-operativity $\eta$ as described in the main text. **(A)** Best designs of controller architecture 1 based on the universal bacterial expression resource and the translational controller. **(B)** Best designs of controller architecture 2 based on the fragmented RNA polymerase and the translational controller.

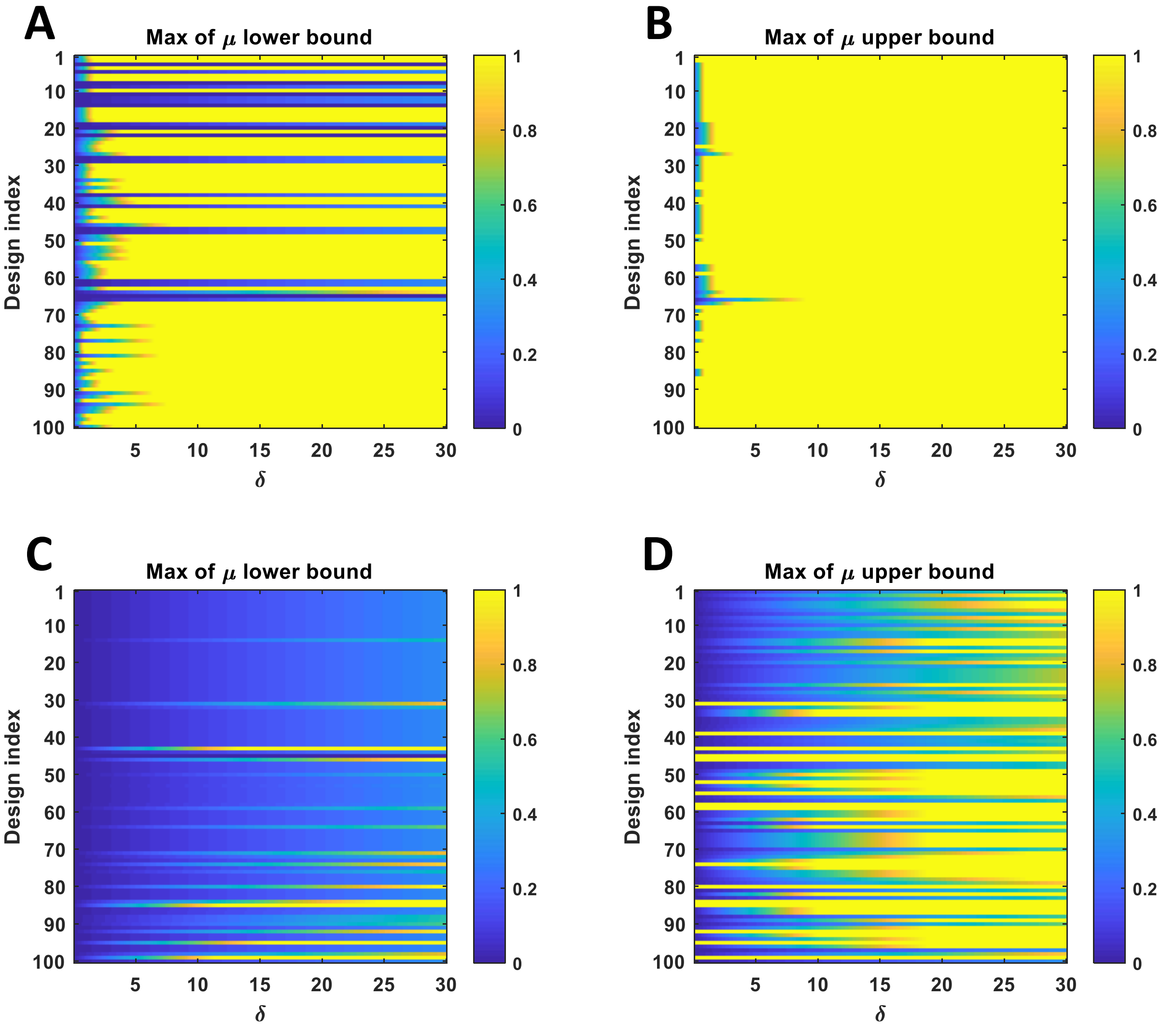


Figure S5: **Extended robustness analysis.** The $\mu$ analysis, as described in the main text, was carried out for all the controllers designed. The maximum of the $\mu$ lower and upper bound estimates where identified. Note that values greater than 1 are cut off at 1 for ease of plotting. If $\mu$ lower bound is greater than 1 then there exists a perturbation of size $\delta$ which will destabilise the system. If $\mu$ lower bound is greater than 1 then stability cannot be guaranteed. **(A)** Maximum value of the $\mu$ lower bound for UBER-OR controllers. **(B)** Maximum value of the $\mu$ upper bound for UBER-OR controllers. **(C)** Maximum value of the $\mu$ lower bound for FRAG-OR controllers. **(D)** Maximum value of the $\mu$ upper bound for FRAG-OR controllers.

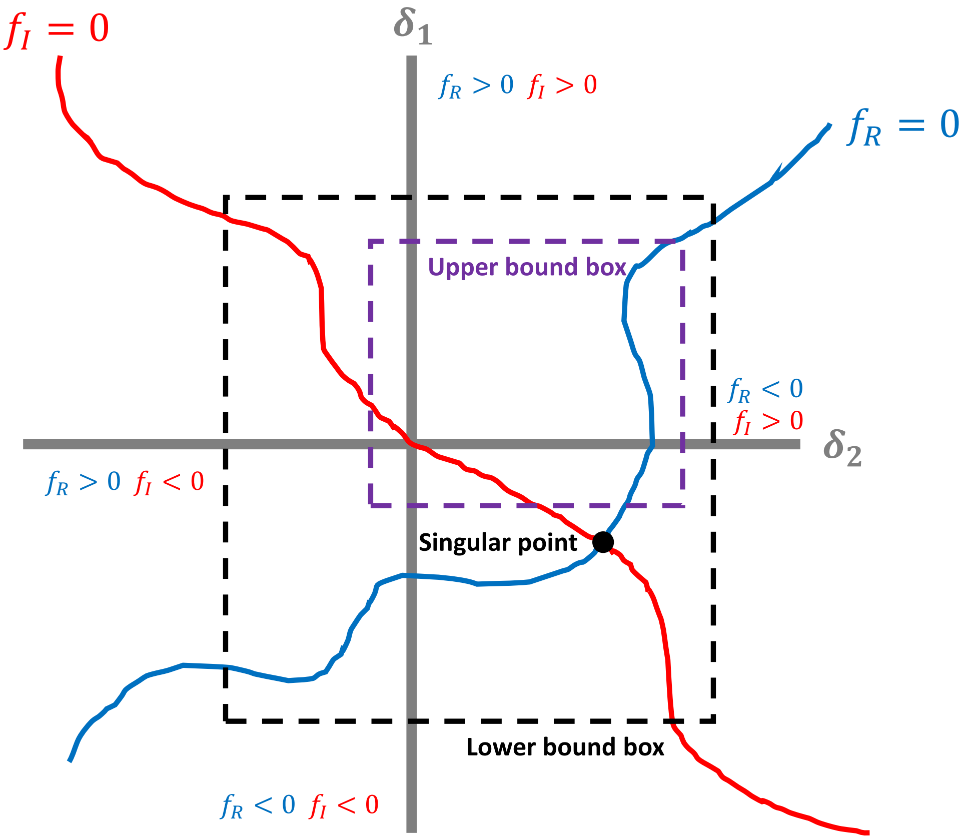


Figure S6: $\mu$ **analysis algorithm.** An example of the $\mu$ analysis algorithm in a simple two dimensional uncertainty space. The two $f_R$ and $f_I$ manifolds stretch across the surface of uncertainty space. The intersection point, labelled singular point, is the true value of $\mu$. The intersection of the two manifolds creates four possible combinations of the signs of $f_R$ and $f_I$. By evaluating the sign combinations covered by two boxes in uncertainty space estimates for $\mu$ can be found. Two boxes are projected onto the uncertainty space and their sizes adjusted until one contains all four $s$ points (this gives the lower bound – i.e. $\mu$ must be within the box) and one which much contains only three of the $s$ points (this gives the upper bound – i.e. $\mu$ cannot be within this box).
